# Optimization of antifungal edible pregelatinized potato starch-based coating formulations by response surface methodology to extend postharvest life of ‘Orri’ mandarins

**DOI:** 10.1101/2020.12.02.408054

**Authors:** Lourdes Soto-Muñoz, Lluís Palou, Maricruz Argente-Sanchis, Miguel Angel Ramos-López, María B. Pérez-Gago

## Abstract

Antifungal composite edible coatings (ECs) formulated with pregelatinized potato starch (PPS, 1.0-2.0 % w/w) as biopolymer, glyceryl monostearate (GMS, 0.5-1.5 %, w/w) as hydrophobe, glycerol (Gly, 0.5-1.5 %, w/w) as plasticizer, and sodium benzoate (SB, 2 % w/w) as antifungal agent were optimized using the Box–Behnken response surface methodology to extend the postharvest life of Orri’ mandarins. The second order polynomial models satisfactorily fitted the experimental data, with high values of the coefficient of determination for the different variables (R^2^>0.91). The individual linear effect of GMS concentration was significant in all the responses evaluated, whereas PPS only affected emulsion viscosity, fruit tacking, and weight loss of coated mandarins. Gly only affected acetaldehyde content in the juice of coated mandarins when interacted with PPS and in the quadratic effect. The optimum concentrations of PPS, GMS, and Gly for the starch-based EC based on maximum fruit quality and required emulsion properties were predicted to be 2.0, 0.5 and 1.0 % (w/w), respectively. The optimal EC reduced weight loss of mandarins and created a modified atmosphere within the fruit without negatively affecting the overall acceptability of the fruit. On the other hand, the optimized antifungal EC containing SB significantly reduced postharvest green and blue molds and sour rot on mandarins artificially inoculated with the pathogens *Penicillium digitatum, Penicillium italicum* and *Geotrichum citri-aurantii,* respectively. Therefore, the optimized antifungal EC showed potential to control the main postharvest diseases and maintain the overall quality of ‘Orri’ mandarins and could be a suitable alternative to commercial citrus waxes formulated with conventional chemical fungicides.

## 1. Introduction

The key goals of postharvest technology for citrus and other fresh fruits are to: *(i)* maintain the quality of the fruit for market requirements, *(ii)* extend their commercial life to achieve a wider distribution of the product, and *(iii)* reduce the losses between harvest and consumption (Zacarías et al., 2020). Several factors contribute to the deterioration of fruit quality and shelf life during postharvest handling. The most important are intrinsic factors related to physiological and biochemical changes that take place during fruit senescence and are related to dehydration of the fruit and increased respiration and ethylene production (Kyriacou and Rouphael, 2018). In addition, there are external factors that contribute to quality deterioration. Among them, infections by postharvest fungal pathogens are considered as the principal cause of citrus wastage, causing large economic losses (Smilanick et al., 2020; Zacarías et al., 2020). Green mold, blue mold, and sour rot, caused by *Penicillium digitatum* (Pers.: Fr.) Sacc., *Penicillium italicum* Wehmer, and *Geotrichum citri-aurantii* (Ferraris) E.E. Butler, respectively, are among the main citrus postharvest diseases worldwide (Smilanick et al., 2020). Currently, the application of synthetic chemical fungicides such as imazalil, thiabendazole, and pyrimethanil, among others, is a common practice to control Penicillium decay, but these fungicides are not effective against *G. citri-aurantii.* Since the relatively recent definitive withdrawal in the European Union (EU) of the active ingredients guazatine (in 2011) and propiconazole (in 2018), there are no authorized fungicide products available for sour rot control in EU citrus producing countries, which is a serious problem for the export sector. This restriction, together with the increasing strong consumer demand for pesticide-free fruits, has encouraged the search for effective and environmentally friendly alternatives to conventional fungicides to control postharvest decay of citrus fruit (Palou et al., 2008, 2016; Soto-Muñoz et al., 2020).

Nowadays, a very active research field is the replacement of commercial citrus waxes, which are often a vehicle for the application of synthetic fungicides, for natural edible coatings (ECs) with antifungal properties (Valencia-Chamorro et al., 2011b). The antifungal activity of such ECs is mainly accomplished through the incorporation to the formulations of non-polluting antifungal ingredients such as plant extracts (Nair et al., 2018), essential oils (Sapper et al., 2019), food additives or low toxicity compounds classified as generally recognized as safe (GRAS) (Guimarães et al., 2019), and microbial antagonists as biocontrol agents (Marín et al., 2017). Among them, GRAS salts show some important advantages such as their high solubility in water, easy availability, and relatively low cost (Palou, 2018). Some previous studies by our group have demonstrated that among GRAS salts, dips in aqueous solutions of sodium benzoate (SB), at concentrations as low as 2-3 % (w/v), have a significant curative activity against citrus green and blue molds (Palou et al., 2002; Montesinos-Herrero et al., 2016) and also sour rot (Soto-Muñoz et al., 2020). Therefore, SB could be an adequate antifungal agent to be incorporated into EC formulations in order to replace the polluting commercial citrus waxes amended with fungicides and maintain the physicochemical and sensory quality of citrus fruits.

Antifungal ECs, in addition to providing antifungal activity, can play an important role in extending the shelf life of fresh fruits by reducing their metabolic activity and maintaining their physicochemical quality (Valencia-Chamorro et al., 2011b; Palou et al., 2015). ECs are typically applied in liquid form to fruits using immersion or spray atomization techniques, leaving a uniform thin layer on the fruit surface. These films create a semi-permeable barrier against gases and water vapor, decreasing the respiration rate and moisture loss, thus contributing to maintain the weight, firmness, and other quality attributes of coated fruit during storage (Sapper and Chiralt, 2018). Thus, for example, composite ECs based on hydroxypropyl methylcellulose (HPMC)-lipid formulated with various food preservatives with antifungal properties to control Alternaria black spot and gray mold in cherry tomatoes (Fagundes et al., 2015), brown rot in plums (Gunaydin et al., 2017), or Diplodia stem-end rot and Penicillium molds in citrus (Guimarães et al., 2019; Valencia-Chamorro et al., 2011a) were reported to reduce respiration rate, ethylene production, weight loss, and fruit firmness during prolonged cold storage. In these research works, the effectiveness and stability of antifungal ECs depended on their compositions and characteristics, and the incorporation of antifungal food additives or GRAS salts greatly modified their barrier and mechanical properties, which made it necessary to optimize the emulsion formulations.

An appropriate approach to optimize coating formulations is the use of Response Surface Methodology (RSM). This methodology permits the optimization of the composition of films and coatings, showing the interactions among variables, with reduced experimental runs (Thakur et al., 2017). RSM has been used in some studies to evaluate the effect of main matrix components (i.e., polysaccharides, proteins, and lipids) and minor components (i.e., plasticizers and emulsifiers) of ECs on fruit quality (Lin et al., 2018). However, the optimization of coating formulations containing food additives with antimicrobial activity has been mainly evaluated *in vitro* using stand-alone films (Mustapha et al., 2019).

Among many polysaccharides studied to develop ECs, potato starch is an excellent film-former, biodegradable, biocompatible, easily available, and low cost compound (Sapper and Chiralt, 2018). In addition, starch films are odorless, tasteless, transparent, and with very low oxygen permeability, which makes this biopolymer a good candidate to coat fresh fruits and vegetables because the created modified atmosphere greatly contributes to an extension of the produce postharvest life (Sánchez-González et al., 2015). Nevertheless, starch-based films and coatings typically have hydrophilic character and poor mechanical properties, for which reason their combination with lipids, minor components such as plasticizers and emulsifiers, or other polysaccharides is required to improve moisture retention, flexibility, extensibility, and/or stability of films and coatings (Sapper and Chiralt, 2018). On the other hand, starch-based coatings are suitable for gaining new functionalities through the incorporation into the formulations of additional ingredients. For example, starch ECs formulated with active antifungal ingredients such as essential oils (Sapper et al., 2019), natamycin-cyclodextrin complex (Yang et al., 2019), lactic acid bacteria (Marín et al., 2019), and GRAS salts like potassium sorbate (Mehyar et al., 2011) have significantly controlled postharvest decay of guava, apple, persimmon, banana, cherry tomato, grapes, and cucumber, among others. However, no information is available regarding the utilization of food additives or GRAS salts as ingredients of starch-based ECs for the control of major fungal postharvest diseases of citrus fruit. Considering the great impact in coating performance of factors such as coating composition (i.e., type of ingredients and relative content), the objectives of the present study were to (i) develop an antifungal EC formulated with SB as antifungal ingredient and different ratios of pregelatinized potato starch (PPS), lipid, and plasticizer optimized by RSM to maintain the physicochemical and sensory quality of ‘Orri’ mandarins during storage; and (ii) evaluate the effect of the optimized EC on the control of citrus postharvest green mold, blue mold, and sour rot on artificially inoculated mandarins.

## 2. Materials and methods

### 2.1. Fruit

The experiments were carried out with ‘Orri’ hybrid mandarins *(Citrus reticulata* Blanco; Saville et al., 2011) at commercial maturity obtained from commercial orchards in the Valencia area (Spain). Fruit were used the same day or stored up to 1 week at 5 °C and 90 % relative humidity (RH) before use. No commercial postharvest treatments were applied before any experiment. Mandarins were selected for uniformity of size and shape and diseased or damaged fruit were discarded. Selected fruit were disinfected superficially by immersion for 4 min in a 0.5% sodium hypochlorite solution, rinsed with tap water, and allowed to air-dry at room temperature before EC application or fungal inoculation. Fruit were appropriately randomized before each experiment.

### 2.2. Emulsion preparation and fruit coating

Edible composite emulsions were prepared by combining PPS (Quimidroga, S.A., Barcelona, Catalonia, Spain), glyceril monostearate (GMS; Italmatch Chemicals Spa, Barcelona, Catalonia, Spain) as lipidic component, and glycerol (Panreac-Química S.A., Barcelona, Catalonia, Spain) as plasticizer suspended in water. Sunflower lecithin and diacetyl tartaric acid esters of mono-diglycerides (Lasenor S.A., Barcelona, Catalonia, Spain) were incorporated as emulsifiers at a ratio GMS: emulsifier of 2:1 (dry basis (db)). Sodium benzoate (SB) at 2 % (w/v) was incorporated into the formulations as antifungal ingredient. PPS solution (5 %, w/w) was stirred at 65 °C for 30 min and maintained under magnetic stirring at 25 °C overnight. Emulsions were prepared as described by Valencia-Chamorro et al. (2011a). Briefly, an aqueous solution of SB at 10 % (w/w) (Sigma-Aldrich Química S.A., Tres Cantos, Madrid, Spain) was prepared. SB solution, sunflower lecithin, diacetyl tartaric acid esters of mono-diglycerides, and water were added to the PPS solution, GMS, and glycerol, at concentrations according to the Box-Behnken experimental design (Table 1) and heated to 90 °C. Once the compounds were melted, samples were homogenized with an UltraTurrax T25 stirrer (Janke&Kunkel, Staufen, Germany) for 1min at 13,000 rpm followed by 3 min at 22,000 rpm. Emulsions were cooled under agitation to a temperature lower than 25 °C by placing them in an ice-water bath under constant agitation for 25 min. The emulsions were kept overnight at 5 °C before use. A set of 60 mandarins per treatment were coated manually by immersion for 15 s at 20 °C in each EC solution (Table 1). Then, coated mandarins were drained and dried in a heated air-forced tunnel at 45±2 °C for 130 s. Fruit were randomly packed into experimental units and stored for up to 2 weeks at 20 °C.

**Table 1.** Matrix of the Box-Behnken experimental design for the optimization of starch based edible coatings

| Run | Factors (% w/w) |  |  |
| --- | --- | --- | --- |
| | $x_1$: Pregelatinized potato starch (PPS) | $x_2$: Glyceryl monostearate (GMS) | $x_3$: Glycerol (Gly) |
| 1 | 2.0 | 1.0 | 0.5 |
| 2 | 1.5 | 1.0 | 1.0 |
| 3 | 1.5 | 1.0 | 1.0 |
| 4 | 1.5 | 1.5 | 0.5 |
| 5 | 1.0 | 0.5 | 1.0 |
| 6 | 1.0 | 1.0 | 0.5 |
| 7 | 2.0 | 1.5 | 1.0 |
| 8 | 2.0 | 1.0 | 1.5 |
| 9 | 1.5 | 0.5 | 0.5 |
| 10 | 2.0 | 0.5 | 1.0 |
| 11 | 1.5 | 1.0 | 1.0 |
| 12 | 1.5 | 0.5 | 1.5 |
| 13 | 1.0 | 1.5 | 1.0 |
| 14 | 1.0 | 1.0 | 1.5 |
| 15 | 1.5 | 1.5 | 1.5 |

### 2.3. Experimental design: Response surface methodology (RSM)

A three level-three factor Box-Behnken RSM design with three central point replicates was employed for the study. Fifteen different EC formulations with the three independent variables, which comprised PPS (*x*_1_): 1.0-2.0 %, GMS (*x*_2_): 0.5-1.5 % and Gly (*x*_3_): 0.5-1.5 %, were used to get optimal combinations. The levels of these independent variables were selected from preliminary single factor tests (data not shown). The experimental design, regression model, and statistical analysis were performed with JMP Statistical Discovery™ software (version 11.0.0; SAS Institute, Cary, NC, USA). All experimental runs are listed in Table 1. The experimental data obtained from experimental runs were fitted according to the following second-order polynomial equation:

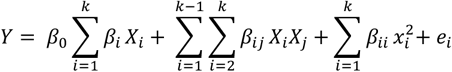

where, *Y* = response or dependent variable; *X_i_* = independent variables; *β*_0_ = intercept; *β_i_, β_ij_, β_ii_* = regression coefficients for intercept, linear, quadratic and interaction terms; k = number of variables.

Contour plots were generated based on the significant effects to visualize the relationship between the response and the independent variables. The adequacy of the polynomial equations was determined by the regression coefficient (R^2^) and the lack-of-fit test. The significance of the model and terms was evaluated by response surface analysis of the experimental design maintaining those terms that showed significant effects at a probability of *p*<0.05. Numerical optimization of the significant variables was performed at least in triplicate to determine the optimal formulation (Rangel-Marrón et al., 2019).

### 2.4. Analytical methods

#### 2.4.1. Emulsion characteristics

##### pH and viscosity

The pH of the emulsions was measured with a pH-meter (Consort C830 multi-parameter analyzer, Turnhout, Belgium) and the viscosity was measured with a viscometer (Visco Star Plus R, Fungilab, S.A., Barcelona, Spain). Measurements were performed in three replicates for each emulsion. Sample viscosities were determined at 20 °C and results were expressed in milipascal-second (mPa-s).

##### Tacking

Five fruit per treatment were manually coated and dried at room temperature (18-23 °C). Coated fruit were placed in a 5-L humidity chamber hermetically closed and incubated at 25 °C and 90 % RH for 24 h. After incubation, fruit were ‘touch-tacky’ and evaluated using a 4-point qualitative scale, were 0=no tacking, 1=low tacking, 2=medium tacking, and 3=high tacking.

#### 2.4.2. Assessment of fruit physicochemical quality

##### Weight loss

Twelve mandarins per treatment were used to measure weight loss by weighing the treated fruit at the beginning of the experiment and at the end of the storage period (2 weeks at 20 °C). Results were presented as the percentage loss of initial weight.

##### Ethanol and acetaldehyde contents

Ethanol and acetaldehyde contents (mg L^-1^) were analyzed from the headspace of 10-mL vials filled with 5 mL of juice from three replicates of 10 mandarins each per treatment according to Valencia-Chamorro et al. (2011b). Samples were analyzed using a GC Thermo Trace (Thermo Fisher Scientific) equipped with an autosampler (Model HS 2000), flame ionization detector (FID), and 1.2 m × 0.32 cm (i.d.) Poropack QS 80/100 column. The injector was set at 175 °C, the column at 150 °C, the detector at 200 °C, and the carrier gas at 28 mL/min.

#### 2.4.3. Assessment of fruit sensory quality

The sensory quality of treated samples was evaluated by 15 trained judges at the end of the 2-week storage period. Five mandarins per treatment were pealed and separated into individual segments. Judges had to taste several segments of each treatment presented in trays labelled with 3-digit random codes. Spring water was provided for palate rinsing between samples. Overall flavor was assessed through a 9-point scale, where 1 to 3 represented a range of non-acceptable quality with the presence of off-flavors, 4 to 6 represented an acceptable quality, and 7 to 9 represented a range of excellent quality. Panelists were also asked to rank visually the treatments from lowest to highest gloss and the summation of rankings were calculated as described by Valencia-Chamorro et al. (2011a).

### 2.5. Effect of coatings on disease development

#### 2.5.1. Fungal inoculum

*P. digitatum* NAV-7, *P. italicum* MAV-1, and *G. citri-aurantii* NAV-1, isolated from rotten citrus fruits found in citrus packinghouses in the Valencia region (Spain), are maintained in the culture collection of postharvest pathogens of the IVIA CTP. They have also been deposited in the Spanish Type Culture Collection (CECT, University of Valencia, Valencia, Spain), where they were assigned the following accession numbers: CECT 21108 to *P. digitatum* NAV-7, CECT 21109 to *P. italicum* MAV-1, and CECT 13166 to *G. citri-aurantii* NAV-1. Conidia from 7-to 14-d-old cultures of these fungi were taken from the PDA (Scharlab S.L., Barcelona, Catalonia, Spain) surface of petri dishes with a sterilized inoculation loop and transferred to a sterile aqueous solution of 0.05 % Tween^®^ 80 (Panreac-Química S.A., Barcelona, Catalonia, Spain). Conidial suspensions were filtered through two layers of cheesecloth. The density of the suspension was measured with a haemocytometer and dilutions with sterile water were done to obtain an exact inoculum density of 10^5^ spores mL^-1^ (*P. digitatum* and *P. italicum)* or 10^7^ arthrospores mL^-1^ (*G. citri-aurantii*). In the case of *G. citri-aurantii*, fresh mandarin juice (10 %), thiabendazole (50 mg L^-1^; Textar^®^60 T, Decco Ibérica PostCosecha, S.A.U., Paterna, Valencia, Spain) and cycloheximide (5 mg L^-1^; Carl Roth GmbH + Co. KG, Karlsruhe, Germany) were added to enhance the infection ability of the arthrospore suspension.

#### 2.5.2. Fruit inoculation, coating application, and disease assessment

Mandarins were artificially inoculated with each pathogen by immersing a stainless-steel rod with a probe tip 1 mm wide and 2 mm in length into the corresponding conidial suspension and wounding each fruit once on the equatorial zone. Mandarins inoculated with *P. digitatum* and *P. italicum* were incubated for 24 h at 20 °C and fruit inoculated with *G. citri-aurantii* were incubated for 24 hat 28 °C. Afterwards, inoculated fruit were coated by immersion (15 s at 20 °C) with the corresponding emulsion, drained, and allowed to air-dry at 20 °C. Treatments were: (*i*) water as uncoated control, (*ii*) the optimal EC formulated without the antifungal GRAS salt (SB), as negative control, (*iii*) 2 % (w/v) SB aqueous solution as positive control, and (*iv*) the optimal antifungal EC (i.e., formulated with 2 % SB). Each treatment was applied to four replicates of five fruit each. All fruit were placed on plastic trays and incubated for 7-days at 20 °C and 90 % RH, at which time disease incidence was assessed as the percentage of decayed fruit and disease severity as the diameter of the lesion (mm).

#### 2.5.3. Statistical analysis

Specific differences between means were determined by Fisher’s protected least significant difference test (LSD, *p*<0.05) applied after an analysis of variance (ANOVA). Analysis was performed using JMP Statistical Discovery™ software (version 11.0.0; SAS Institute). For incidence data, the ANOVA was applied to the arcsine of the square root of the proportion of decayed fruit. Non-transformed means are shown.

## 3. Results and discussion

### 3.1. Experimental design analysis

The emulsion characteristics (pH and viscosity), fruit tacking, and the physicochemical and sensory quality parameters of coated mandarins stored for 14 days at 20 °C (weight loss, ethanol and acetaldehyde content in juice, overall flavor, and visual fruit gloss rank) are shown in Table 2. The experimental data were used to determine the coefficients of the quadratic polynomial equations of the dependent variables. Regression analysis and Pareto ANOVA were conducted to fit the model and determine the statistical significance of the model terms. The coefficient of determination R^2^, adjusted R^2^, the lack of fit, and F-ratio of the model with the corresponding *p*-value for each dependent variable are given in Table 3. R^2^ values ranged from 0.906 to 0.985, indicating an excellent relationship between the model and the experimental data. The lack of fit values were not statistically significant (*p*<0.05), confirming that the model adequately describes the suggested quadratic surface and explains the experimental data. Furthermore, the high F-values of the model (from 5.379 to 37.572) and *p*-values lower than 0.05 imply that the model was significant and reliable to predict the emulsion characteristics and the physicochemical and sensory quality of coated mandarins stored for 2 weeks at 20 °C.

**Table 2.** Experimental data obtained for the response variables studied in antifungal starch-based edible coatings and on coated ‘Orri’ mandarins stored at 20 °C for 2 weeks

| Run | Emulsion characteristics |  |  | Fruit physicochemical and sensory quality |  |  |  |  |
| --- | --- | --- | --- | --- | --- | --- | --- | --- |
| | pH | Viscosity<br>(mPa-s) | Tacking<br>(0-3) | Weight loss<br>(%) | Ethanol<br>(mg L<sup>-1</sup>) | Acetaldehyde<br>(mg L<sup>-1</sup>) | Overall flavor<br>(1-9) | Ranked fruit gloss<br>($\Sigma$ rank) |
| 1 | 5.50 | 175.25 | 2.9 | 3.83 | 927.0 | 9.6 | 4.9 | 59 |
| 2 | 5.47 | 32.71 | 2.0 | 3.77 | 835.0 | 12.2 | 4.9 | 70 |
| 3 | 5.45 | 34.24 | 2.5 | 3.85 | 585.7 | 9.2 | 4.9 | 70 |
| 4 | 5.25 | 138.82 | 2.8 | 4.33 | 1237.3 | 18.9 | 4.9 | 26 |
| 5 | 5.73 | 49.52 | 0.3 | 3.58 | 473.2 | 10.2 | 6.0 | 103 |
| 6 | 5.48 | 35.98 | 0.9 | 4.12 | 802.1 | 12.7 | 5.3 | 80 |
| 7 | 5.36 | 281.67 | 2.7 | 4.09 | 1619.7 | 21.1 | 4.2 | 62 |
| 8 | 5.47 | 139.83 | 2.8 | 3.62 | 745.2 | 13.1 | 4.3 | 93 |
| 9 | 5.67 | 24.08 | 0.6 | 3.67 | 380.6 | 7.8 | 6.0 | 132 |
| 10 | 5.71 | 40.46 | 0.9 | 3.16 | 560.8 | 12.4 | 6.3 | 98 |
| 11 | 5.45 | 24.95 | 2.4 | 3.83 | 978.6 | 13.0 | 5.5 | 57 |
| 12 | 5.71 | 25.10 | 0.2 | 3.45 | 406.9 | 10.0 | 6.1 | 104 |
| 13 | 5.37 | 43.64 | 1.8 | 3.92 | 1587.9 | 22.7 | 4.6 | 31 |
| 14 | 5.45 | 35.24 | 1.0 | 4.25 | 509.3 | 7.9 | 5.6 | 82 |
| 15 | 5.30 | 62.03 | 1.5 | 3.98 | 1047.2 | 15.9 | 4.6 | 49 |

Tacking scale: 0= null, 1=low, 2=medium, 3=high; Overall flavor scale: 1-3=bad, 4-6=acceptable, 7-9=excellent.

**Table 3.** Analysis of variance of emulsion characteristics and physicochemical and sensory quality of coated ‘Orri’ mandarins for determination of model fitting

| Variable | R<sup>2</sup> | Adjusted R<sup>2</sup> | Lack of fit | F ratio of the model | *p* value>F |
| --- | --- | --- | --- | --- | --- |
| pH | 0.983 | 0.952 | 0.0719<sup>ns</sup> | 31.572 | 0.0007* |
| Viscosity | 0.985 | 0.959 | 0.0641<sup>ns</sup> | 37.253 | 0.0005* |
| Tacking | 0.912 | 0.755 | 0.1744<sup>ns</sup> | 5.782 | 0.0339* |
| Weight loss | 0.934 | 0.814 | 0.0651<sup>ns</sup> | 7.816 | 0.0178* |
| Ethanol | 0.929 | 0.801 | 0.6547<sup>ns</sup> | 7.261 | 0.0209* |
| Acetaldehyde | 0.966 | 0.904 | 0.8799<sup>ns</sup> | 15.626 | 0.0037* |
| Global acceptability | 0.906 | 0.738 | 0.4383<sup>ns</sup> | 5.379 | 0.0393* |
| Ranked fruit gloss | 0.919 | 0.774 | 0.1746<sup>ns</sup> | 6.316 | 0.0281* |

\* Significantly different ( $p<0.05$ ).
<sup>ns</sup> Not significantly different ( $p>0.05$ ).

### 3.2. Analysis of response surfaces

#### 3.2.1. Emulsion characteristics

##### pH and viscosity

pH values of the EC emulsions were moderately acid and ranged between 5.25 and 5.73 (Table 2). The R^2^ value of 0.983 is in satisfactory correlation with the adjusted R^2^ value of 0.952, verifying that the model describes 98.3 % of the variation (Table 3). Furthermore, the non-significant lack of fit value (0.0719) indicates the high precision and reliability of the model. Table 4 shows the regression coefficients of the individual linear, quadratic, and interaction terms and their statistical significance for pH. The results showed that the pH was highly influenced by the concentration of GMS, both in the linear and quadratic terms (*p*<0.05), while the content of PPS and glycerol did not affect the pH of the formulations. The pH of the emulsions had a negative correlation with the GMS variable (i.e., increasing the GMS concentration decreased the pH) (Fig. 1a). This may be associated with the acidic nature of GMS. GMS is a mixture of monoglycerides of stearic and palmitic acids together with varying amounts of di- and tri-glycerides and smalls amounts of free stearic, palmitic, and oleic acids, with a pH value lower than 6.0 (Rowe et al., 2009).

**Fig. 1.**
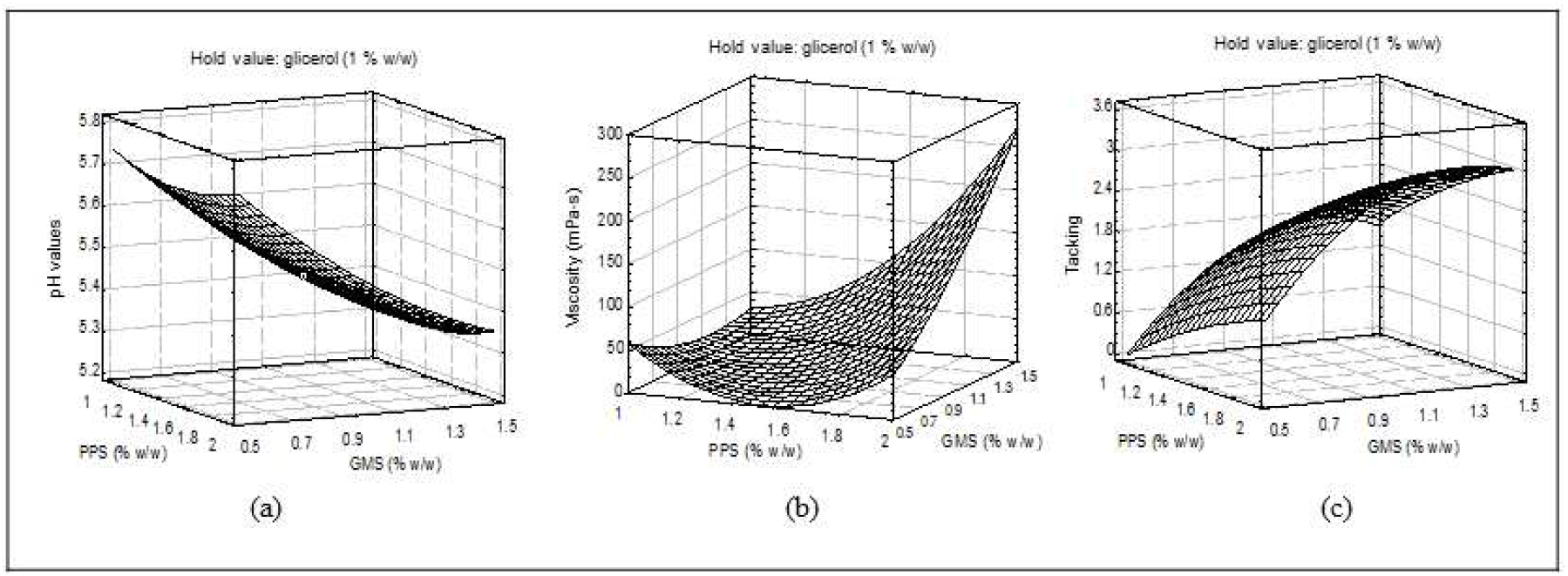
Response surface plots showing the effect of the interactions among process variables on emulsion characteristics: (a) pH, (b) viscosity, and (c) fruit tacking.

**Table 4.** Regression coefficients and significance probability (*p*-value) of experimental data

| Parameters | pH |  | Viscosity (mPa-s) |  | Tacking |  | Weight loss (%) |  | Ethanol (mgL<sup>-1</sup>) |  | Acetaldehyde (mg L<sup>-1</sup>) |  | Global acceptability |  | Ranked fruit gloss |  |
| --- | --- | --- | --- | --- | --- | --- | --- | --- | --- | --- | --- | --- | --- | --- | --- | --- |
|  | Estimate | Prob>\|t\| | Estimate | Prob>\|t\| | Estimate | Prob>\|t\| | Estimate | Prob>\|t\| | Estimate | Prob>\|t\| | Estimate | Prob>\|t\| | Estimate | Prob>\|t\| | Estimate | Prob>\|t\| |
| $\beta_0$ | 5.456 | <0.0001* | 30.6333 | 0.0173 | 2.3 | 0.0004* | 3.814 | <0.0001* | 799.74 | 0.0005* | 11.4667 | <0.0001* | 5.11 | <0.0001* | 65.6666 | 0.0003* |
| $\beta_1$ | 0.001 | 0.919 | 59.104 | <0.0001* | 0.662 | **0.012\*** | -0.1478 | **0.023\*** | 60.04 | 0.380 | 0.34 | 0.535 | -0.2325 | 0.118 | 3.25 | 0.532 |
| $\beta_2$ | -0.193 | <0.0001* | 48.375 | **0.0003\*** | 0.85 | **0.004\*** | 0.3065 | **0.001\*** | 458.831 | **0.0007\*** | 4.802 | **0.0002\*** | -0.7688 | **0.002\*** | -33.625 | **0.001\*** |
| $\beta_3$ | 0.0038 | 0.762 | -13.991 | 0.049 | -0.212 | 0.269 | -0.0828 | 0.139 | -79.806 | 0.257 | -0.265 | 0.626 | -0.0563 | 0.667 | 3.875 | 0.460 |
| $\beta_{12}$ | 0.0025 | 0.886 | 61.773 | **0.0005\*** | 0.075 | 0.769 | 0.1475 | 0.078 | -13.947 | 0.881 | -0.96 | 0.241 | -0.17 | 0.374 | 6.5 | 0.386 |
| $\beta_{13}$ | 0 | 1.000 | -8.67 | 0.304 | -0.05 | 0.844 | -0.0847 | 0.259 | 27.742 | 0.766 | 2.085 | **0.034\*** | -0.22 | 0.263 | 8 | 0.295 |
| $\beta_{23}$ | 0.0025 | 0.886 | -19.4525 | 0.050 | -0.225 | 0.394 | -0.035 | 0.621 | -54.09 | 0.567 | -1.305 | 0.130 | -0.1025 | 0.583 | 12.75 | 0.122 |
| $\beta_{11}$ | 0.0392 | 0.072 | 53.6283 | **0.001\*** | -0.125 | 0.640 | -0.0142 | 0.845 | 119.275 | 0.251 | 1.4142 | 0.119 | -0.1187 | 0.542 | 4.291667 | 0.574 |
| $\beta_{22}$ | 0.0467 | **0.043\*** | 19.5608 | 0.056 | -0.75 | **0.031\*** | -0.1125 | 0.166 | 141.373 | 0.184 | 3.724 | **0.004\*** | 0.2787 | 0.185 | 3.54166 | 0.640 |
| $\beta_{33}$ | -0.0208 | 0.281 | 12.3133 | 0.179 | -0.275 | 0.324 | 0.1542 | 0.077 | -173.12 | 0.118 | -2.051 | **0.041\*** | 0.0137 | 0.942 | 8.54166 | 0.284 |

\* Significantly different ( $p<0.05$ ); $\beta_0$ : intercept; $\beta_1, \beta_2$ and $\beta_3$ : linear regression coefficients for pregelatinized potato starch (PPS), glyceryl
monostearate (GMS) and glycerol (Gly); $\beta_{12}, \beta_{13}$ and $\beta_{23}$ : regression coefficients for interactions between PPS × GMS, PPS × Gly and GMS × Gly;
$\beta_{11}, \beta_{22}$ and $\beta_{33}$ : quadratic regression coefficients for PPS × PPS, GMS × GMS and Gly × Gly.

In general, the influence of pH on *in vivo* fungal growth is complex and dependent upon the ionization of acids or bases in the medium in which the fungus resides. In the case of many citrus fruits, including mandarins, pH of the rind tissues, where infections by wound pathogens such as *P. digitatum, P. italicum*, and *G. citri-aurantii* initiate and develop, is between 5 and 5.5 (Palou et al., 2002), which is very similar to the pH of most of the emulsion combinations tested in this work (Table 2). Therefore, coating application to the mandarin surface would only change minimally the pH of the rind. Some studies have reported that the undissociated molecule of benzoic acid is responsible for the antimicrobial activity of the salt SB (Valencia-Chamorro et al., 2011b). Thus, the maximum antimicrobial effect of SB is at a pH value of 4.20, which is the pKa of benzoic acid in solution and corresponds to the highest concentration this form can reach. However, the pH of SB aqueous solutions at concentrations around 2 % is slightly higher than neutral (around 7.5) (Palou et al., 2002), so theoretically, and from this point of view, these ECs formulated with SB would be more effective to inhibit citrus wound pathogens than SB solutions applied as a stand-alone treatment.

Emulsion viscosity is an important parameter to be considered in the coating process. For instance, applying the coating using spray atomization techniques requires a lower viscosity than the immersion technique, although both techniques require a low viscosity at the commercial scale. In the current study, the viscosity of the EC emulsions for all the experimental combinations varied between 24.95 and 281.67 mPa-s, and approximately a viscosity value of 40 mPa-s was considered as the maximum limit to ensure a correct application and to minimize any possibility of plugging the spray nozzle during hypothetical application in citrus processing lines in commercial packinghouses. The high values of R^2^ (0.985) and adjusted R^2^ (0.959) indicate a good fit of the model, with 98.5 % of the observed variation in viscosity explained by it (Table 2). PPS and GMS showed a significant positive linear effect on viscosity. Likewise, the interaction PPS × GMS and the quadratic term of PPS also showed a positive effect on emulsion viscosity (*p*<0.05) (Table 4). As clearly shown in Fig. 1b, the emulsion viscosity increased with an increase in GMS concentration and there was a strong interaction with the PPS content, so that an increase in the PPS/GMS content implied a significant increase in viscosity to reach values close to 300 mPa-s. In general, potato starch has been reported to form pastes with a higher viscosity than other commercial starches (Singh et al., 2003). This has been attributed to the high content of phosphate ester groups that covalently bond to amylopectin molecules, contributing to enhance the starch viscosity (Singh et al., 2004). These rheological properties of starch can be altered by the presence of polar lipids, such as monoglycerides (Singh et al., 2003). In our work, the effect of GMS on the emulsion viscosity depended on the starch concentration. At low concentration of starch, the addition of the lipid probably retarded the swelling and solubility of the starch and hence decreased the emulsion viscosity, whereas, at high concentrations of starch, the addition of the GMS resulted in higher viscosity, probably due to a more rigid starch granule by the action of the lipids (Deffenbaugh, 2019).

##### Tacking

As shown in Table 2, fruit tacking values ranged from 0.2 (no tacking) to 2.9 (high tacking). The high R^2^ value indicated that 91.2 % of the total variation was explained by the model (Table 3). Results in Table 4 show that PPS and GMS had a significant positive linear effect on tackiness of coated mandarins, while the quadratic term of GMS had a significant negative effect (*p*<0.05). The variable Gly and the rest of interactions between independent variables had no significant effect on tacking. In general, when ECs are used as surface coatings for fruits, they should not be tacky after drying and maintain this property during storage to prevent them from sticking to packaging materials or fingers, especially after exposure to humid environments. According to our results, PPS in combination with GMS at concentrations above 1 % caused the coatings to be tacky (i.e., tacking > 1) (Fig. 1c). This could be attributed to the high-water absorption capacity and swelling power of starch granules when exposed to high RH conditions (Kim et al., 2015), and to the high viscosity of the emulsions containing high concentrations of GMS that could have modified the amount of coating adhered to the fruit surface, increasing the absorption of water into the film matrix (Fernández-Cervera et al., 2004).

#### 3.2.2. Physicochemical quality parameters of mandarin fruits

##### Weight loss

Weight loss of coated mandarins after 2 weeks of storage at 20 °C varied from 3.16 to 4.33 % (Table 2). As shown in the response surface plot, weight loss of coated mandarins decreased as the PPS concentration increased and, contrastingly, increased with increasing GMS concentrations (Fig. 2a). In general, starch-based films have a high water vapor permeability derived from the hydrophilicity of starch. Therefore, the addition of hydrophobic compounds such as waxes, oils, mono and di-glycerides, fatty acids, and other lipids are incorporated into the formulations to improve the water vapor barrier properties (González-Soto et al., 2019; Sapper and Chiralt, 2018). However, issues such as lipid type, volume fraction, and polymorphic phase affect the barrier properties of emulsion films (Pérez-Gago and Rhim, 2014). Furthermore, in addition to these factors, coating performance is affected by surface interactions between the coating-forming emulsion and the fruit peel, and a good surface wettability, proper adhesion, and durability during storage are required properties (Sapper and Chiralt, 2018). Fruit skin morphology (i.e., thickness and type of cuticle, presence of hairs, number of stomata, lenticels, and even cracks in the lenticels) and coating physical properties, such as surface tension and viscosity, strongly influence mass transfer of the coated fruit (Navarro-Tarazaga et al., 2011). In our work, at low GMS content (0.5 %), an increase in PPS concentration significantly reduced the weight loss of mandarins to reach the lowest value (3.2 %). This could be related to an increase in the coating thickness as the EC solid content increased, improving the coverage of the mandarins. However, weight loss of mandarins increased to 4.10 % as GMS content increased, independently of the PPS content. In spite of a continuous reduction in water vapor permeability in emulsion films as the hydrophobic components increased, some studies have found a critical volume fraction beyond which the barrier properties of the emulsion films did not improve or even got worse. For instance, emulsion films of corn starch and sunflower oil showed a minimum permeability with about 2.2 g L^-1^ lipid content, beyond which the water barrier decreased (Garcia et al., 2000). Similarly, Petersson and Standing (2005) reported that the addition of an ester of monoglycerides increased the water vapor permeability of a potato starch-based film. This was attributed to the monoglyceride modifying the starch polymer matrix by breaking hydrogen bonds or by hindering chain-chain interactions and thereby giving an increased diffusion of water vapor in the film. Similar results have also been reported for coated fruit such as plums, where an increase in beeswax content incorporated into HPMC increased weight loss of coated fruit. This was attributed to coating brittleness, which might have featured some discontinuities, cracks or holes, reducing the water barrier of the coating (Navarro-Tarazaga et al., 2011). On the other hand, monoglycerides such as GMS can also act as surfactants in contact with other lipids, which could partially remove the natural waxes present in the fruit cuticle, resulting in an increase in weight loss (Harker and Ferguson, 1991).

**Fig. 2.**
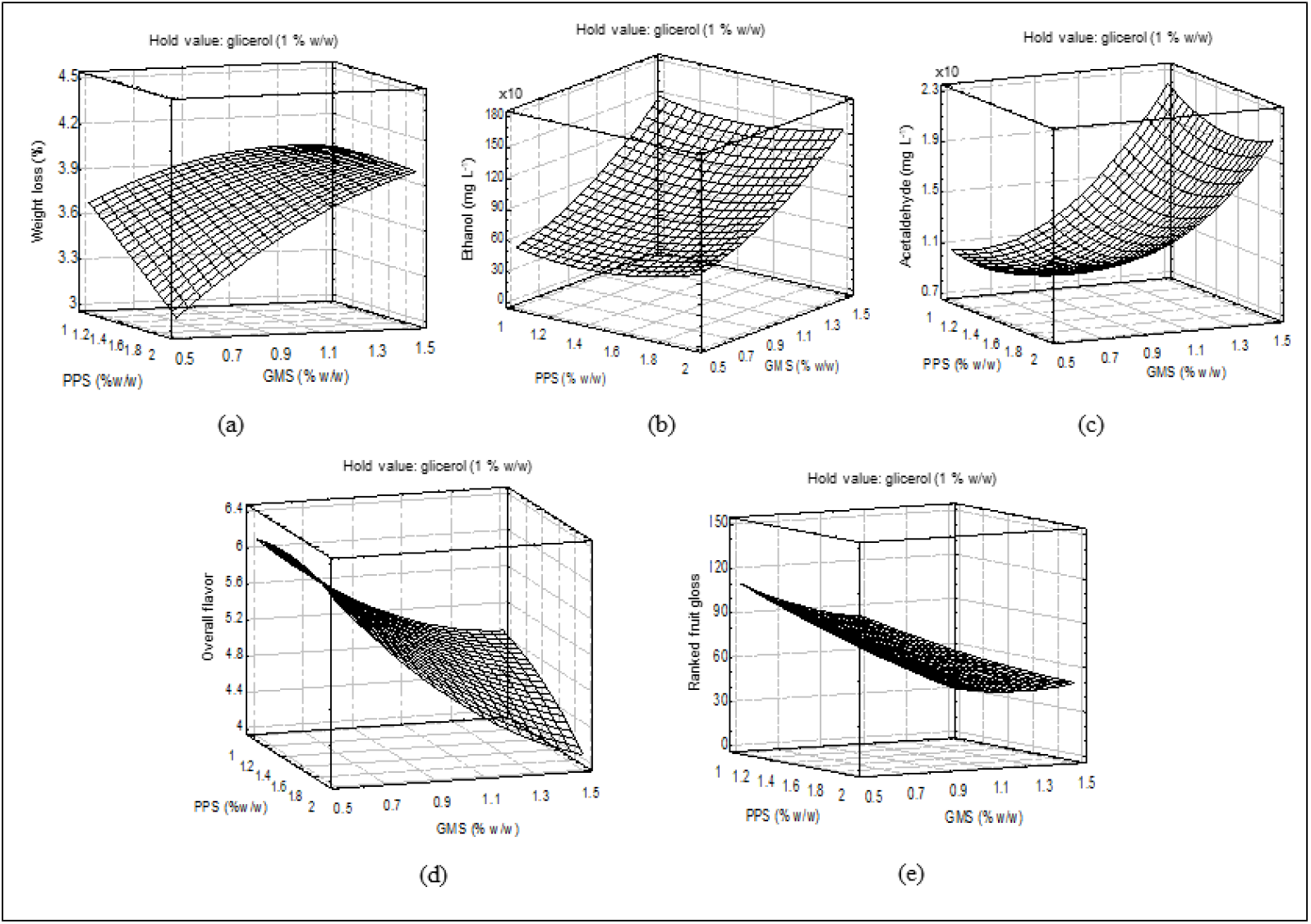
Response surface plots showing the effect of the interactions among process variables on physicochemical and sensory quality of coated ‘Orri’ mandarins stored at 20 °C for 2 weeks: (a) weight loss, (b) ethanol content, (c) acetaldehyde content, (d) overall flavor, and (e) visual fruit gloss rank.

##### Ethanol and acetaldehyde contents

As shown in Table 2, the content of ethanol and acetaldehyde in the juice of coated mandarins ranged from 380.6 to 1619.7 mg L^-1^ and 7.8 to 22.7 mg L^-1^, respectively, after 2 weeks of storage at 20 °C. The R^2^ values obtained for ethanol and acetaldehyde content were 0.929 and 0.966, respectively, indicating a goof fit of the model (Table 3). Among the independent variables, GMS showed a significant positive linear effect for both volatiles (*p*<0.05), with a significant increase as GMS concentration increased (Fig. 2b,c). This increase in the content of internal volatiles associated with anaerobic conditions indicates an increase in the gas barrier properties of the ECs due to GMS incorporation into the starch polymer matrix, whereas the PPS content had a minimum effect. This result seems to contradict some authors that have reported that the incorporation of hydrophobic components to hydrophilic polymer matrixes, such as starch or cellulose derivatives, increased the oxygen permeability due to the higher chemical affinity and solubility of hydrophobic components compared to hydrocolloid polymers (Cazón et al., 2017). However, these results could be related to the viscosity of the EC formulations. Many works have reported a direct relationship between the internal gas modification of coated fruit and coating thickness, which depends basically on the solid content and the viscosity of the coating formulations (Contreras-Oliva et al., 2012; Yousuf and Qadri, 2020). On the other hand, Ben-Yehoshua et al. (1985) concluded that the resistance of citrus fruits to mass transport of water vapor and gases occurred by different mechanisms. Thus, while water moves preferentially through the cuticle, gases diffuse mainly through the stomatal openings. Therefore, fruit coating can partially or completely plug stomatal pores, greatly reducing gas exchange, whereas the presence of cracks or lack of uniformity in the coating layer might not improve the final moisture barrier. This fact could explain the results observed in the present work regarding weight loss and volatile content of ‘Orri’ mandarins coated with ECs containing high content of GMS.

In addition to GMS, the acetaldehyde content was also positively affected by the interaction PPS × Gly and the quadratic term of GMS, and negatively affected by the quadratic term of Gly (Table 4). The effect of Gly modifying the acetaldehyde content could be related to the plasticizer effect reducing the interactions between the starch-starch molecules, resulting in a loser polymer network, as described by Bertuzzi et al. (2007).

#### 3.2.3. Sensory quality of mandarin fruits

Fitting the model for the variables overall flavor and ranked visual fruit gloss gave R^2^ values of 0.906 and 0.919, respectively, and the values of lack of fit, F, and *p* verified the reliability of the model to predict them (Table 3). Both responses were negatively affected by the linear term of GMS (*p*<0.05) (Table 4). Fig. 2d,e represents the response surface plots for overall flavor and the summation of fruit gloss ranks of coated mandarins as a function of the content of PPS and GMS in the ECs. An increase in the GMS concentration resulted in a decrease in the overall flavor, from 6.3 to 4.2 after 2 weeks of storage at 20 °C. These values were in the range of acceptability (6-4), although the response of the ECs with high GMS content was in the lower limit due to a slight appreciation of off-flavor by the panelists associated with an increase of ethanol and acetaldehyde contents in the juice. Several research works show that the contribution to off-flavor of volatile content depends on citrus cultivar. In general, ethanol levels above 2,000 mg L^-1^ have been associated with off-flavors in ‘Valencia’ oranges and ‘Oronules’ mandarins, while other mandarin cultivars such as ‘Ortanique’ and ‘Fortune’ had an ethanol threshold for off-flavor detections above 3,000 mg L^-1^ (Contreras-Oliva et al., 2012; Perez-Gago et al., 2002). In the present study, the maximum ethanol value was 1,620 mg L^-1^ in mandarins coated with ECs formulated with high PPS and GMS content, and these corresponded with the lower overall flavor, indicating the high susceptibility of ‘Orri’ mandarins to off-flavor development.

On the other hand, high-ranking gloss values were scored when fruit were coated with formulations containing low GME amounts and the values declined when GMS content in the EC formulations increased (Fig. 2e). Our results are in agreement with those obtained by Pérez-Gago et al. (2002), where increasing the lipid content from 20 to 60 % (db) in HPMC-based emulsion coatings decreased the gloss of coated mandarins. This was attributed to the translucent character of the emulsion formulations that increased with lipid content and/or the coating thickness, which increased with the viscosity of the emulsion. Similarly, it has been reported for different biopolymer-lipid emulsion films that the incorporation of lipids significantly decreased the film gloss compared to the pure hydrocolloid film and the particle size of the dispersed phase influenced gloss values (Trezza and Krochta, 2000).

### 3.3. Optimization and validation of model

The desirability function was applied for simultaneous optimization of the multiple responses. PPS, GMS, and Gly contents of the antifungal EC tailored for coating of ‘Orri’ mandarins will be considered optimum when the emulsion complies with the following characteristics: maximum pH value, viscosity below 40 mPa-s, and null fruit tacking, and the quality of coated mandarins after storage complies with the smallest possible values of weight loss, ethanol and acetaldehyde content, and the highest sensory quality (overall flavor and gloss). For these conditions, the optimum level for the different variables was achieved with 2.0 % PPS, 0.5 % GMS, and 1.0 % Gly (w/w), with an overall desirability of 0.86.

The adequacy of the models was experimentally validated with the above-mentioned optimal conditions. Table 5 shows the results of triplicate experiments compared with the predicted values of the responses. The results demonstrated that experimental values were reasonably close to predicted values. Except for acetaldehyde content, there were no significant differences between predicted and experimental values (*p*>0.05), indicating the suitability of the methodology used for the optimization of the variables and the accuracy of the reduced models fitted by RSM.

**Table 5.** Validation of predicted values for emulsion characteristics and quality parameters of ‘Orri’ mandarins coated with an optimized potato starch-based antifungal edible coating*

| Response variable | Predicted value | Experimental value<sup>a</sup> |
| --- | --- | --- |
| pH value | 5.73a | 5.73±0.01a |
| Viscosity (mPa-s) | 40.0a | 44.7±4.3a |
| Tacking | 1.0a | 0.4±0.5a |
| Weight loss (%) | 3.2a | 3.13±0.8a |
| Ethanol content (mg L<sup>-1</sup>) | 634.2a | 774.5±206.10a |
| Acetaldehyde content (mg L<sup>-1</sup>) | 12.19a | 19.4±4.2b |
| Global acceptability | 6.0a | 5.2±1.5a |

\*Composition of the optimal coating was: pregelatinized potato starch: 2 % (w/w); glyceryl monostearate: 0.5 % (w/w); and glycerol: 1% (w/w). All the coatings contained 2% sodium benzoate (w/w) as antifungal agent.
<sup>a</sup>Values are means ± standard deviations.
Values in the same row with the same letter are not significantly different ( $p<0.05$ ).

### 3.4. Effect of the optimized coating on disease development

The optimized EC formulation was evaluated to control disease caused by *P. digitatum, P. italicum*, and *G. citri-aurantii* on mandarins artificially inoculated, coated 24 h later, and incubated for 1 week at 20 °C (green and blue molds) or 28 °C (sour rot). The optimized EC formulated without the antifungal ingredient SB and immersion in 2 % SB aqueous solution (positive control) were also assayed. The incidence and severity of Penicillium molds and sour rot on uncoated mandarins (treated with water) and those coated with the EC without antifungal agent were not significantly different, which indicated that the EC without SB did not exhibit *in vivo* antifungal activity against any of the three fungi (Fig. 3). In contrast, the optimized antifungal EC (formulated with 2 % SB) significantly reduced the incidence and severity of green and blue molds and sour rot on ‘Orri’ mandarins. The effectiveness of the optimized antifungal EC was significantly higher to control sour rot than green and blue molds, and no significant differences in incidence were found between the two Penicillium molds. The optimized EC reduced the incidence of green and blue molds by about 30 % and the incidence of sour rot by 55 %. (Fig. 3a). Likewise, a similar pattern was observed for disease severity. Thus, the optimized antifungal EC reduced green and blue molds severity by 52-64 % and sour rot severity by 66 % with respect to control fruit after 7 days of incubation (Fig. 3b). On the other hand, the effectiveness of the optimized antifungal EC to reduce the incidence of blue mold and sour rot was similar to that of the 2 % SB aqueous treatment. However, the antifungal coating was significantly less effective in reducing the incidence of green mold (about 30 % reduction) than the 2 % SB solution (about 70 % reduction), and the same behavior was observed for disease severity.

**Fig. 3.**
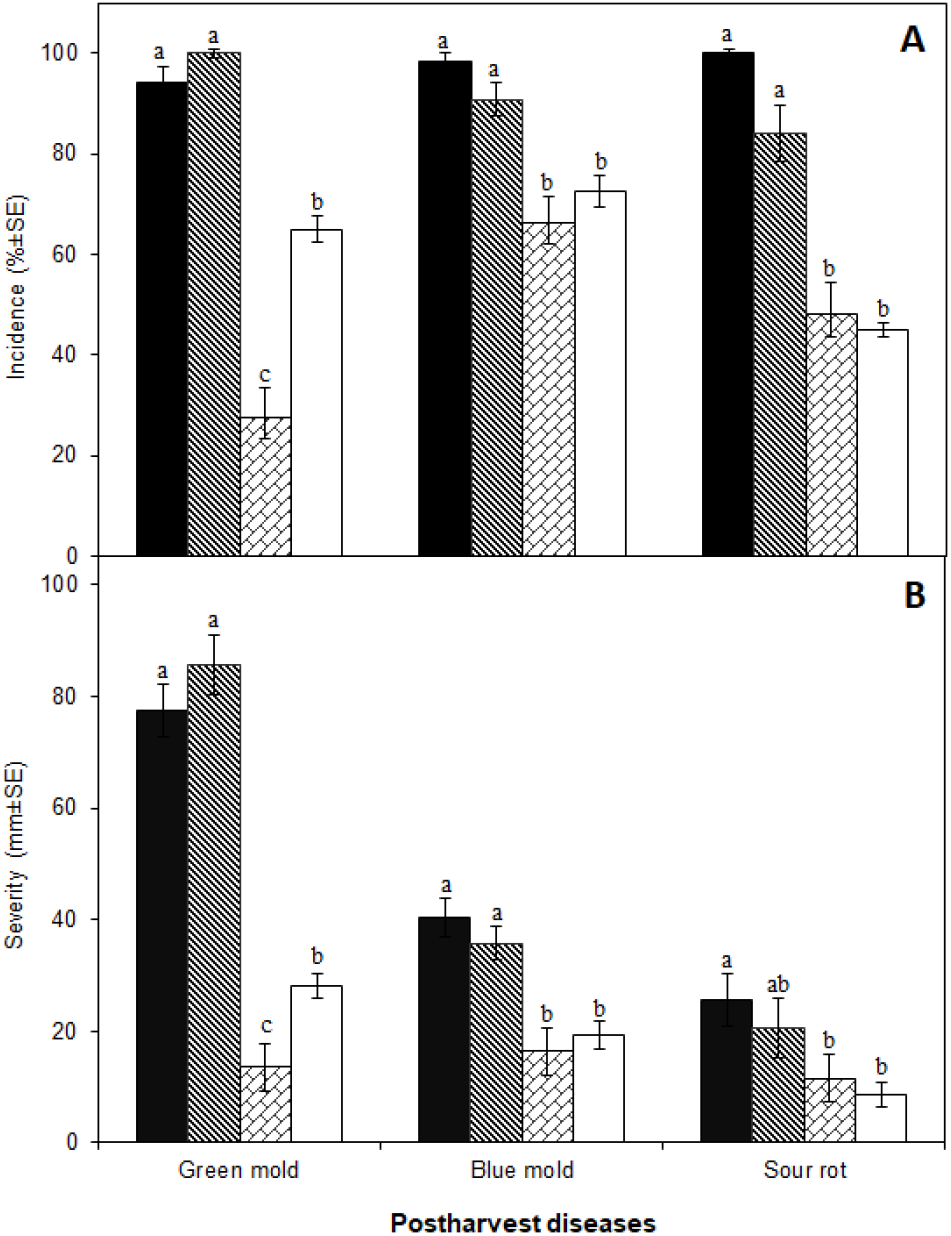
Disease incidence (A) and severity (B) of green mold, blue mold, and sour rot on ‘Orri’ mandarins artificially inoculated with *Penicillium digitatum*, *Penicillium italicum*, and *Geotrichum citri-aurantii,* respectively, treated 24 h later with water (Control;■), edible coating (EC) formulated without the antifungal ingredient sodium benzoate (SB) 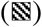, 2 % SB aqueous solution 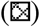, or optimized antifungal EC (□), and incubated for 7 days at 20 °C (green and blue molds) or 28 °C (sour rot) and 90 % RH. Values are means of 4 replicates of 5 fruit each per treatment and all assays were carried out in triplicate. For each disease, mean values with different letters indicate significant differences among treatments according to Fisher’s protected LSD test (*p*<0.05). For disease incidence, the ANOVA was applied to the arcsine-transformed values. Non-transformed means are shown.

These results confirm that SB was the coating ingredient responsible for the antifungal activity of the optimized EC in this study and are consistent with other studies that demonstrated the efficacy of SB applied as aqueous solutions to control postharvest diseases caused by *Penicillium* spp. and *G. citri-aurantii* on a variety of citrus species and cultivars (El-Mougy et al., 2008; Montesinos-Herrero et al., 2016; Palou et al., 2002; Muñoz-Soto et al., 2020). Moreover, the effectiveness of HPMC-lipid ECs containing SB was highly effective to control Penicillium decay on ‘Clemenules’ mandarins (Valencia-Chamorro et al., 2011a) and ‘Valencia’ oranges (Valencia-Chamorro et al., 2009). A particularly interesting result was the high curative activity that the optimal EC showed to control citrus sour rot. Very scarce information is available on the reduction of disease caused by *G. citri-aurantii* by antifungal ECs. Taking into account that the postharvest fungicides currently registered in the EU for citrus fruits are not effective to control sour rot, the use of ECs formulated with antifungal agents could be an effective and safe strategy to be implemented commercially in citrus packinghouses as a replacement for the conventional waxes used for physiological preservation.

The similar effect of the optimal antifungal EC and the SB aqueous solution to control decay, at least in the case of blue mold and sour rot, suggests that the emulsion formulation and properties allow the release of SB from the coating to the fruit surface and its diffusion into the pathogen infection courts (fruit rind wounds). This is a very important factor to consider since the formulation of ECs with the addition of antifungal ingredients such as food additives, essential oils, or other GRAS compounds may allow an increase of the activity of the antifungal agent by regulating its temporal and spatial release or facilitating its continuous and effective contact with the target pathogen. Other substantial advantages of antifungal ECs compared to the use of antifungal agents as stand-alone treatments are related to their mode of application. Since ECs can be applied with standard waxing equipment already present in most citrus packinghouses, there is no need for the acquisition of new equipment technologies or the allocation of new facilities that might be needed for implementation of novel gaseous or aqueous treatments. Furthermore, coating application may considerably reduce the risks of phytotoxicity or induction of adverse flavors or odors associated with postharvest treatment with GRAS substances such as plant extracts, essential oils, and other volatile compounds (Chen et al., 1996; Palou et al., 2016).

In the case of green mold, the present results showed significant differences in both disease incidence and severity between fruit treated with the optimal EC and aqueous SB. As just discussed above, factors other than salt release and diffusion from the EC matrix should explain why aqueous SB controlled green mold more effectively than the antifungal EC, because both treatments similarly controlled blue mold and sour rot and the EC formulated without SB showed no control ability. pH can obviously play a role, but as previously discussed in subsection 3.2.1, the optimal EC (with a pH of 5.73; Table 5) should exert higher antifungal activity in the citrus rind (pH of 5-5.5) than the SB salt aqueous solution (pH of 7.5) and this is not the case. However, as pointed out by Zhang et al. (2013), factors additional to the proportion of the acidic form present in the solution may have an impact on the ability to control citrus green mold. In their work, these authors found that neutral or alkaline pH conditions disrupted the transcription factor *PdpacC* and the polygalacturonase gene *Pdpg2,* resulting in impaired mycelial growth and attenuated virulence of *P. digitatum*.

## 4. Conclusion

A new antifungal starch-based EC with potential for use during postharvest handling of ‘Orri’ mandarins was successfully developed using a three level Box-Behnken response surface design. The optimized antifungal EC contained PPS, GMS, glycerol, sunflower lecithin, diacetyl tartaric acid esters of mono-diglycerides, and SB as antifungal ingredient, and met specific requirements for proper commercial application: (i) the emulsion was stable and had low viscosity to allow its application in citrus processing lines, (ii) the formulation covered the fruit surface adequately and the fruit was not sticky after drying, (iii) the EC effectively reduced weight loss and, although increased the ethanol and acetaldehyde content in the juice, did not induce off-flavor in coated mandarins after storage, (iv) the EC maintained the gloss and natural appearance of coated mandarins, and (v) the EC significantly reduced green and blue molds and sour rot caused by *P. digitatum, P. italicum*, and *G. citri-aurantii*, respectively, on artificially inoculated ‘Orri’ mandarins. Overall, this starch-based coating containing SB could be a promising treatment to maintain quality, reduce decay, and extend postharvest life of mandarins, and thus effectively substitute the use of conventional waxes amended with synthetic chemical fungicides. Its potential commercial use as part of integrated disease management programs would be especially suitable for the control of sour rot, since in important citrus producing areas such as the EU, the conventional chemical fungicides currently approved for postharvest use on citrus fruits are not effective against this disease. Further research should be conducted to evaluate the effect of the application of this antifungal EC at a commercial scale and on other citrus species and cultivars during cold storage and shelf life.

## Acknowledgements

This work was partially funded by the IVIA (Project No. 51910) and the European Commission (FEDER Program). The authors gratefully acknowledge Fontestad S.A. (Montcada, Valencia) for providing fruit and technical assistance and Lasenor S.A. (Barcelona, Catalonia) for providing the lecithin and acid esters of mono-diglycerides. Lourdes Soto-Muñoz’s postdoctoral program is supported by a scholarship by the Mexican National Council of Science and Technology (CONACYT-160058, México).

